# A conserved triple arginine motif in OMA-1 is required for RNA-binding activity and embryo viability

**DOI:** 10.1101/2025.05.09.653132

**Authors:** Asli Ertekin, Sharon T. Noronha, Christable Darko, Francesca Massi, Sean P. Ryder

**Author notes:** Co-corresponding authors; 364 Plantation Street LRB-925, Worcester, MA, 01605, USA; 364 Plantation Street LRB-906, Worcester, MA, 01605, USA. These authors contributed equally to this work (Asli Ertekin, Sharon T. Noronha).

## Abstract

Sexually reproducing organisms make haploid gametes—oocytes and spermatocytes—that combine during fertilization to make an embryo. While both gametes contain similar DNA content, oocytes contain the bulk of the cytoplasm including maternally supplied mRNAs and proteins required prior to zygotic gene activation. RNA-binding proteins are key regulators of these maternal transcripts. In *Caenorhabditis elegans*, the tandem zinc finger proteins OMA-1 and OMA-2 are required for fertilization. Here, we show that OMA-1 RNA-binding activity requires a short basic region immediately upstream from the canonical tandem zinc finger domain. Mutation of this region in animals produces a phenotype distinct from a genetic null. Oocytes can be fertilized, but fail to form an intact chitin egg-shell, frequently break in utero, and arrest prior to morphogenesis. Our results identify a critical region outside of the canonical RNA-binding domain required for both RNA-binding activity as well as revealing a new role for OMA-1 during the oocyte-to-embryo transition.

## INTRODUCTION

Fertilization is the complex and highly coordinated process of cellular fusion between two haploid gametes to form a new diploid zygote. The female gamete, or oocyte, must undergo a maturation process to be fertilized [1]. Maturation steps include the completion of meiosis and breakdown of the nuclear envelope, among others. These steps are conserved throughout the plant and animal kingdoms and are found in all sexually reproducing species.

In the model organism *Caenorhabditis elegans*, immature oocytes are arrested in prophase of diakinesis I [2]. Maturation begins when the most proximal oocyte receives a maturation signal from a secreted sperm-produced protein called the major sperm protein (MSP) [3, 4]. This signal initiates nuclear envelope breakdown, rearrangement of the cortical cytoplasm, and completion of meiosis I, which occurs concurrently with penetration of the male gamete (spermatocyte) [5]. Following fertilization, meiosis II begins, polar bodies are extruded, a hard chitin shell is formed, and the pro-nuclei fuse to create the new zygote’s nucleus [5]. This process, known as the oocyte-to-embryo transition [6], occurs once every twenty minutes in this species.

Two redundant genes (*oma-1* or *oma-2*) are necessary for oocyte maturation in *C. elegans* [7]. Either is sufficient, but when both are lost, oocytes fail to mature, stacking up in the germline. No embryos are produced. OMA-1 and OMA-2 proteins both contain a CCCH-type Tandem Zinc Finger (TZF) domain like their mammalian homolog tristetraprolin (TTP, ZFP36) [8, 9]. The TZF domain is a canonical RNA-binding domain that binds directly to RNA sequences through base and backbone specific contacts [10-14]. TTP promotes mRNA turnover through binding to AU-rich elements in the 3’-untranslated region (UTR) of its target mRNAs [15, 16]. We previously showed that OMA-1 also binds with high affinity to RNA sequences containing UAU and UAA elements [17]. However, OMA-1 exhibits strong positive cooperativity in RNA-recognition, a feature not observed for other members of the TZF family.

OMA-1 and OMA-2 are thought to bind and negatively regulate many maternal transcripts keeping them silent throughout the oocyte-to-embryo transition [7, 17, 18]. Accordingly, OMA-1 has been shown to directly associate with ∼1500 transcripts in RNA-immunoprecipitation (RIP) studies [18], and *oma-1/oma-2* RNAi leads to the deregulation of many 3’UTR reporter transgenes, including *zif-1, mei-1, glp-1, atg-4*.*2, set-6, cul-4, ets-4*, and *mex-3* [19-21]. In another role, OMA-1 and OMA-2 are thought to repress transcription of zygotic genes in the early embryo through association with TAF-4 [22]. The precise mechanism, or mechanisms, by which OMA-1 and OMA-2 coordinates the oocyte-to-embryo transition is not known.

In this study, we set out to define the molecular basis for OMA-1’s RNA-binding affinity and cooperativity. Surprisingly, our results reveal that the canonical TZF domain is not sufficient to support high affinity RNA-binding. Instead, RNA-recognition requires the TZF domain and an upstream thirteen amino acid arginine rich motif. Mutating three adjacent arginine residues to alanine reduces the affinity by more than 3.5-fold. Engineering the same RRR to AAA substitution into the endogenous *oma-1* locus in *Caenorhabditis elegans* causes a fully penetrant embryonic lethality phenotype in the absence of *oma-2*. Unlike a genetic null or TZF domain deletion, the RRR to AAA mutation produces oocytes that can be fertilized. However, the embryos fail to hatch and are characterized by weak eggshells that frequently break, polynucleated embryonic cells, and embryonic arrest prior to morphogenesis. Together, our results identify a critical region of *oma-1* outside of the TZF domain essential for RNA binding and reveal a new role for OMA-1 in promoting eggshell synthesis during oocyte-to-embryo transition.

## RESULTS

### The oma-1 TZF domain is not sufficient for RNA binding activity

CCCH-type tandem zinc finger proteins are composed of two highly conserved zinc finger motifs connected by a flexible linker [8]. This structured domain is flanked by disordered low complexity N- and C-terminal domains, both of which exhibit little sequence conservation (**Fig. 1A, Supplemental Figure 1**). There have been several quantitative studies on the RNA binding activity of members of this protein family [11-14, 16, 17, 23, 24]. The isolated TZF domains from human proteins TIS11D (residues 152-220) and TTP (residues 102-170) bind to AU-rich RNA recognition sequences with an apparent equilibrium dissociation constant (K_D,app_) of 9 ± 2, and 9 ± 1 nM, respectively [12]. Similar studies on the *C. elegans* proteins MEX-5 and POS-1 also showed that the isolated TZF domains (residues 268-346 for MEX-5 and 97-173 for POS-1) bind to their target RNA with K_D,app_ of 16 ± 1 nM and 58 ± 2 nM, respectively [13, 14].

**Figure 1.**
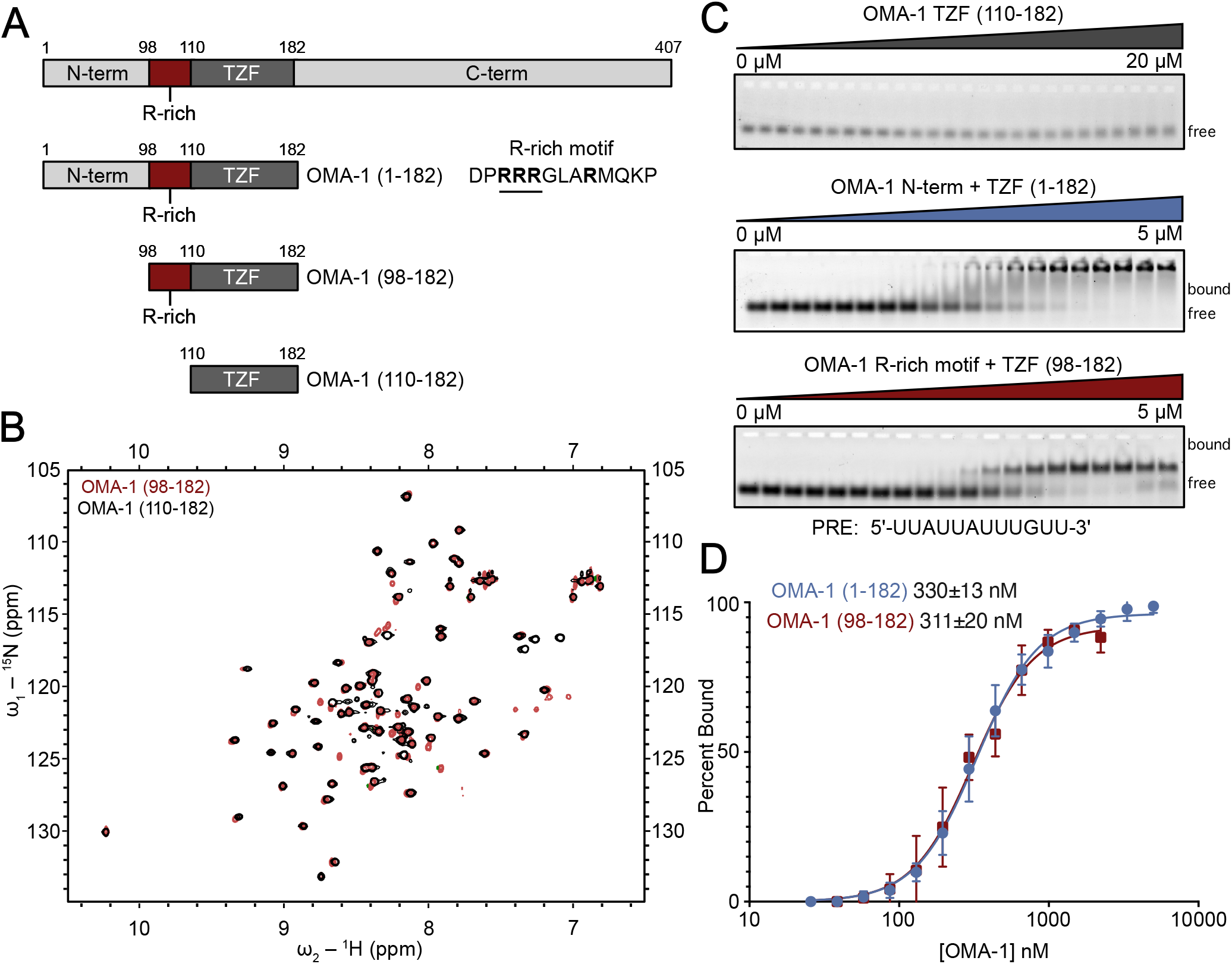
An arginine-rich region in OMA-1 is necessary for RNA-binding activity. **A**. Schematic of the OMA-1 protein constructs used in this work. The amino acid regions are noted. The TZF domain and R-rich motif are labeled. **B**. HSQC spectrum of OMA-1(92-182) (red) and OMA-1(110-182) (black) showing that both protein variants are folded and adopt similar conformations. **C**. Representative EMSA gels with all three protein variants. Bound and free RNA species are labeled. The RNA sequence used is shown below the gels. **D**. Binding curves used to determine the apparent equilibrium dissociation constants (shown) for OMA-1(92-182) (red) and OMA-1(1-182) (blue).

To better understand how the OMA-1 TZF domain structure and dynamics contribute to its RNA-binding activity, we expressed this domain (residues 110-182) in *E. coli* and purified for use in NMR spectroscopy and RNA binding studies. Previous studies showed that maltose binding protein (MBP)-tagged OMA-1(1-182) bound a UAU/UAA rich selected aptamer sequence with high affinity (K_D,app_ = 15 ± 1 nM) and high positive cooperativity (n_H_ = 3.5) [17]. The protein also bound with moderate affinity and reduced cooperativity to a shorter sequence from the 3’ UTR of *glp-1* mRNA termed the POS-1 recognition element (PRE, K_D,app_ = 112 ± 47 nM, n_H_ = 1.3) [17, 24].

The well-dispersed peaks observed in the ^15^N-^1^H-HSQC spectrum indicate that both zinc fingers of the OMA-1 TZF domain are folded in the absence of RNA (**Fig. 1B**). To determine whether the OMA-1 TZF domain is sufficient for RNA binding, we performed electrophoretic mobility shift assays (EMSA) with purified OMA-1 (110-182) and the *glp-1* 3’UTR sequence described above [25]. Interestingly, we did not observe any binding (**Fig. 1C**), suggesting that, unlike other members of this RNA-binding protein family, the TZF domain of OMA-1 alone is not sufficient for RNA binding.

### An arginine rich region is necessary for high affinity binding

Arginine residues play a crucial role in nucleic acid recognition in a variety of protein-RNA and protein DNA complexes [26-29]. The arginine side chain enables a multimodal interaction pattern, including πstacking, π-cation interactions, electrostatic interactions, and H-bond formation with the nucleic acid. Arginine side chains contribute to both sequence and structure specific interactions as well as non-specific backbone interactions between proteins and RNA [26, 30]. A 2001 study analyzing 32 protein-RNA complexes observed that arginine is involved in 26% of all hydrogen bond contacts and 14% of all Van der Waals contacts [26].

The OMA-1 TZF domain is preceded by an arginine-rich region, a thirteen amino acid stretch containing four arginine residues (**Fig. 1A**). Given the importance of arginine residues in RNA recognition, we hypothesized that this region may be required for the RNA-binding activity of OMA-1. To test this hypothesis, we purified two protein variants—one containing the entire N-terminal region and TZF domain (1-182) and another containing the arginine-rich region and the TZF domain (98-182)—and tested their ability to bind the PRE sequence using EMSA (**Fig. 1C**). We fitted the resulting binding curves using the Hill equation to determine the apparent K_D_ and the Hill coefficient (n) (**Fig. 1D**). For the first construct, OMA-1(1-182), the K_D,app_ is 330 ± 13 nM and the n is 2.1 ± 0.1. For OMA-1(98-182), the K_D,app_ is 311 ± 20 nM and the n is 2.2 ± 0.2. The results show that inclusion of the short arginine-rich region preceding the TZF domain is sufficient to fully support OMA-1 RNA binding activity. Superimposition of the ^15^N-^1^H-HSQC spectra for OMA-1(98-182) and OMA-1(110-182) shows that the structure of the TZF domain is unchanged between the two constructs (**Fig. 1B**), indicating that the arginine rich region does not cause large structural perturbations to the TZF domain.

### The arginine rich motif contributes to cooperative RNA-binding activity

To test whether the arginine residues preceding the TZF contribute to the binding affinity, we mutated three adjacent arginine residues—R100, R101, R102—into alanine or lysine residues in the Hexa-his-MBP-OMA-1(1-182) construct, OMA-1(1-182)-AAA and OMA-1(1-182)-KKK, respectively (**Fig. 2A**). The RNA-binding activity of MBP-tagged OMA-1(1-182)-WT, OMA-1(1-182)-AAA and OMA-1(1-182)-KKK were tested against several RNA sequences that were previously studied, including native sequences and an *in vitro* selected aptamer sequence (**Fig. 2B-D**) [17].

**Figure 2.**
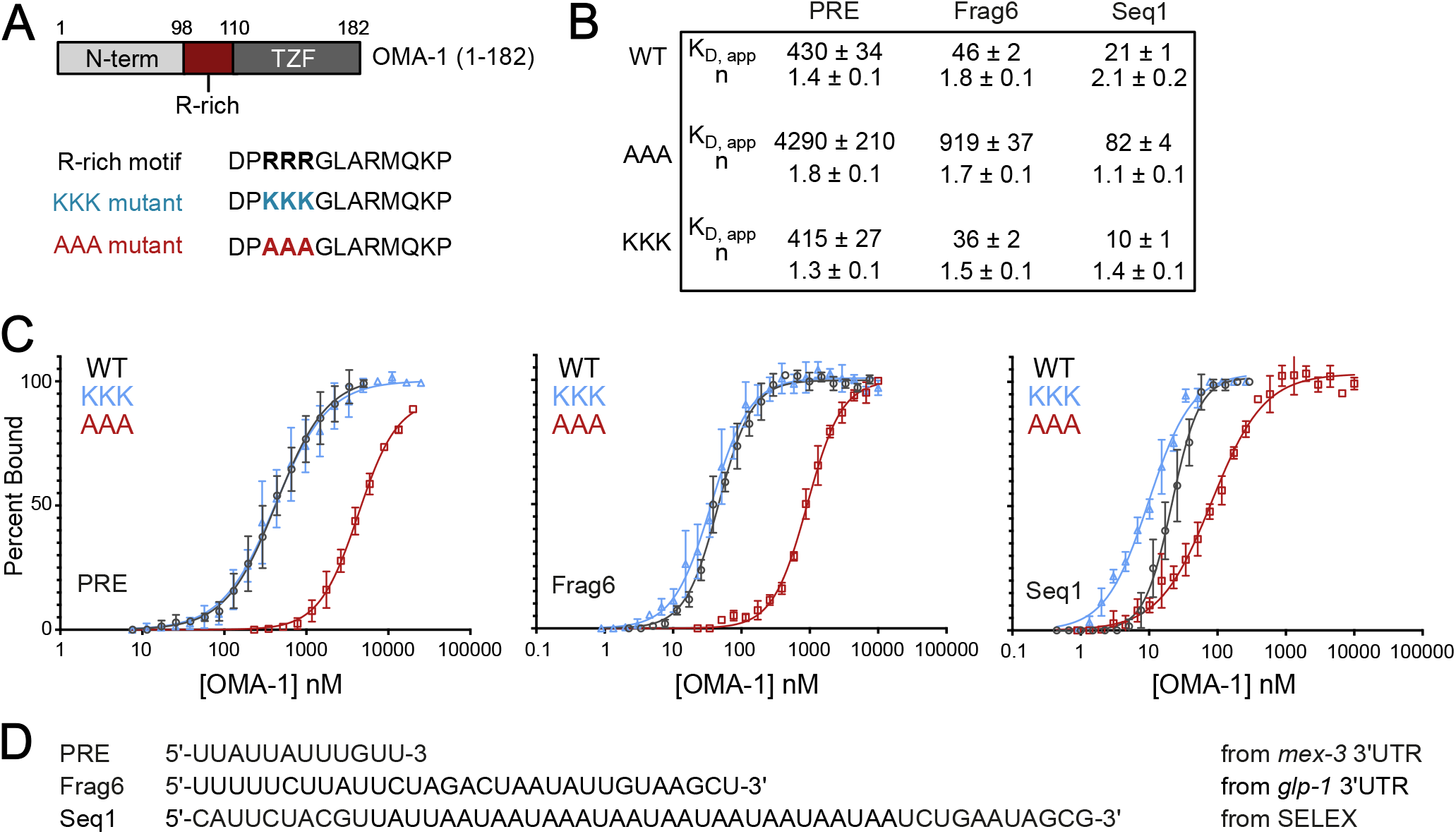
A triple arginine motif outside of the TZF contributes to high affinity RNA-binding. **A**. Schematic of the OMA-1 variant (1-182) and the R-rich peptide. The KKK and AAA mutant variants are shown. **B**. The apparent equilibrium dissociation constants (K_D,app_) and Hill Coefficients (n) of the binding curves shown in (**C**) are presented for three different RNA sequences. Curves for WT are in black, the KKK mutant are blue, and the AAA mutant are red. **D**. The sequences used for the binding experiments are shown, and the source of the sequence from is shown.

EMSA binding curves for the three proteins with three different RNA sequences are shown in **Figure 2C**. In addition to the PRE sequence, two other sequences, seq1 and frag6, were selected from an earlier study [17]. Seq1 was recovered from a systematic evolution of ligands by exponential enrichment (SELEX) protocol to identify the binding motif of OMA-1, and frag6 corresponds to a region in the 3’-UTR of the OMA-1 target *glp-1*. The results show that mutating the triple arginine motif to lysine residues has little effect on binding of OMA-1 to all sequences. In contrast, we observe a significant weakening of the RNA-binding activity for the OMA-1(1-182)-AAA construct. The apparent K_D_ for the PRE sequence is increased by 10 fold in the mutant (K_D,app,AAA_ = 4290 ± 210 nM) compared to the WT (K_D,app,WT_ = 430 ± 34 nM) and triple lysine mutant (K_D,app,KKK_ 415 ± 27 nM). Using frag6, the triple alanine mutant shows a ∼20-fold decrease in binding (K_D,app, AAA_ = 919 ± 37 nM) compared to the WT (K_D,app,WT_ = 46 ± 2 nM) and the triple lysine mutant (K_D,app,KKK_ = 36 ± 2 nM). Using the selected aptamer (seq1), we measured a K_D,app_ of 82 ± 4 nM (OMA-1(1-182)-AAA), 21 ± 1 nM (OMA-1(1-182)-WT), and 10 ± 1 nM (OMA-1(1-182)-KKK).These results show that across multiple RNA targets that bind with a range of affinities, the OMA-1(1-182)-AAA mutant shows significantly weaker binding (4-fold to 20-fold) compared to wild-type (RRR) and triple lysine mutations, suggesting a role for electrostatic forces in the interaction.

We also observe that binding cooperativity is reduced in OMA-1(1-182)-AAA compared to OMA-1(1-182)-WT or -KKK when binding to seq1, which contains multiple OMA-1 binding sites (**Fig. 2B-D**). The mechanism underlying these differences in cooperativity remains unclear. A positive correlation between the reported Hill coefficient for several different RNA sequences and their free energy of folding suggests that the secondary structure of the target RNA may play a role in the cooperativity of binding (**Supplemental Figure 2**). The positively charged residues, RRR/KKK, may interact with the bases to disturb the secondary structure of the RNA, exposing the other binding sites, which would have been protected otherwise.

### A model strain to evaluate strong oma-1 alleles

Given these results, we were interested to test the impact of *oma-1* RNA-binding domain mutations *in vivo*. Because *oma-1* and *oma-2* are redundant, it is relatively straightforward to recover strong loss-of-function alleles of each gene. However, to evaluate phenotypes caused by these alleles, it is necessary to cross in a null allele of the other gene. Because inactivation of both genes causes sterility, we must perform this cross each time we set up a new experiment to evaluate the phenotype. To get around this bottleneck, we generated a strain (WRM66, **Fig. 3A–B**) with the following three modifications. First, GFP was knocked into the endogenous *oma-1* locus enabling visualization of the OMA-1 expression pattern. Second, an auxin-inducible degron (AID) tag was knocked into the endogenous *oma-2* locus, enabling rapid depletion using the auxin degron system [31]. Third, a Tir1::mRuby transgene was integrated into the genome using mosSCI [32]. This transgene is expressed in the germline and early embryos.

**Figure 3.**
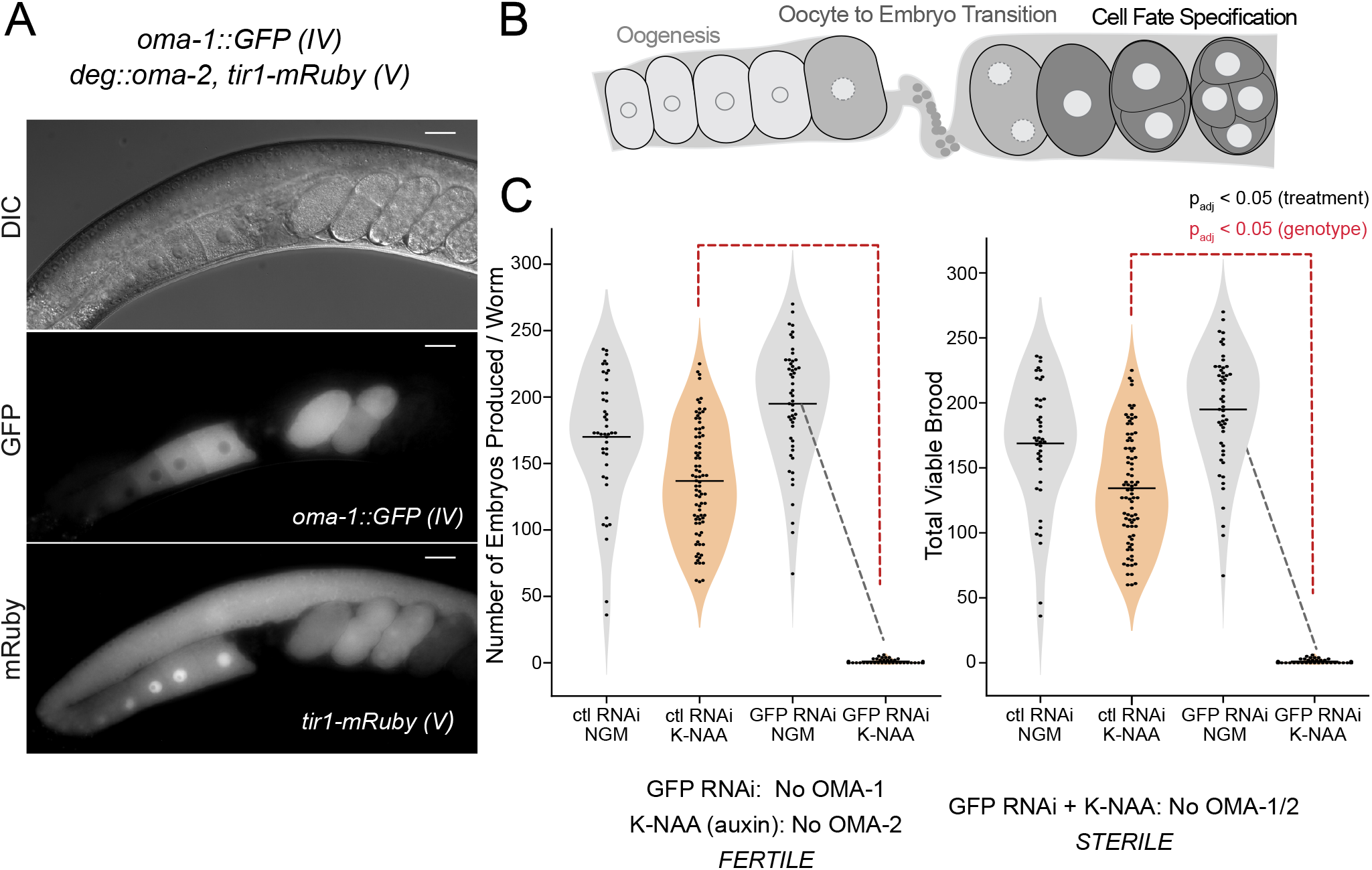
A strain to recover and evaluate strong alleles of *oma-1*. **A**. The genotype of the background strain used in this work (WRM66), along with a representative image showing the oocytes and embryos (DIC), the pattern of OMA-1::GFP expression (GFP), and the pattern of expression of a Tir1::mRuby transgene (mRuby) expressed in the germline and early embryos needed for auxin-mediated depletion of OMA-2 protein. The scalre bar represents 20 microns. **B**. Schematic of ovulation in *C. elegans*, including late-stage oogenesis, sperm, the oocyte-to-embryo transition (fertilization), and early embryogenesis. **C**. A violin plot displaying the total and viable brood per adult hermaphrodite is presented in the presence and absence of control or GFP-targeting RNAi when cultured on standard nematode growth media (NGM, gray) or NGM supplemented with K-NAA (a soluble auxin salt, orange). Each dot represents the brood from a single animal. The solid black bar represents the mean of all animals measured. The dashed lines indicate statistical significance (p_adj_ < 0.05) between culture conditions (black) or between RNAi treatments (red) in a one-way ANOVA with Bonferroni correction.

This strain is expected to have the following characteristics. First, when *oma-1* is inactivated, either by mutation or by GFP-targeted RNAi, the animals remain fertile in the absence of auxin but are sterile in the presence of auxin. Second, if *oma-1* is functional, animals remain fertile in the presence or absence of auxin. Third, if *oma-1* mutations are hypomorphic, the animals display reduced fertility on auxin and normal fertility in the absence of auxin. To test these predictions, we measured the brood size of animals treated with GFP-targeting RNAi or a non-targeting control RNAi in the presence and absence of K-NAA, an auxin analog (**Fig. 3C**). Animals treated with GFP-targeting RNAi but not K-NAA produced 195 ± 45 viable progeny per adult hermaphrodite (N=51). Animals treated with GFP RNAi and K-NAA produced 1 ± 1.6 embryos per adult hermaphrodite (N=45, p_adj_ < 0.05, one way ANOVA with Bonferroni correction). By comparison, animals treated with control RNAi produced viable progeny both in the presence and absence of K-NAA (brood_CTL RNAi, -KNAA_ = 169 ± 48; brood_CTL RNAi, +KNAA_ = 134 ± 43, N = 43 and 83 respectively). The results show that sterility is observed only when both OMA-1 (GFP RNAi) and OMA-2 (K-NAA) are knocked down. The data confirm the robustness of the strain for evaluation of *oma-1::gfp* alleles without the need for genetic crosses.

### OMA-1 RNA-binding activity is essential to fertility

Next, we prepared two *oma-1* mutants in this strain background using CRISPR-Cas9 genome editing. In the first, we made a precise in-frame deletion of the *oma-1* TZF domain (amino acids 110–182, COP2772: *oma-1::GFP(ΔTZF)*). In the second, we mutated *oma-1* R100-R102 codons to alanine codons (WRM92: *oma-1::GFP(AAA)*). Both mutants produce visible OMA-1::GFP protein at the expected location in late stage oocytes and early embryos at similar abundance, suggesting neither mutation disrupts protein expression or localization (**Fig. 4A**).

**Figure 4.**
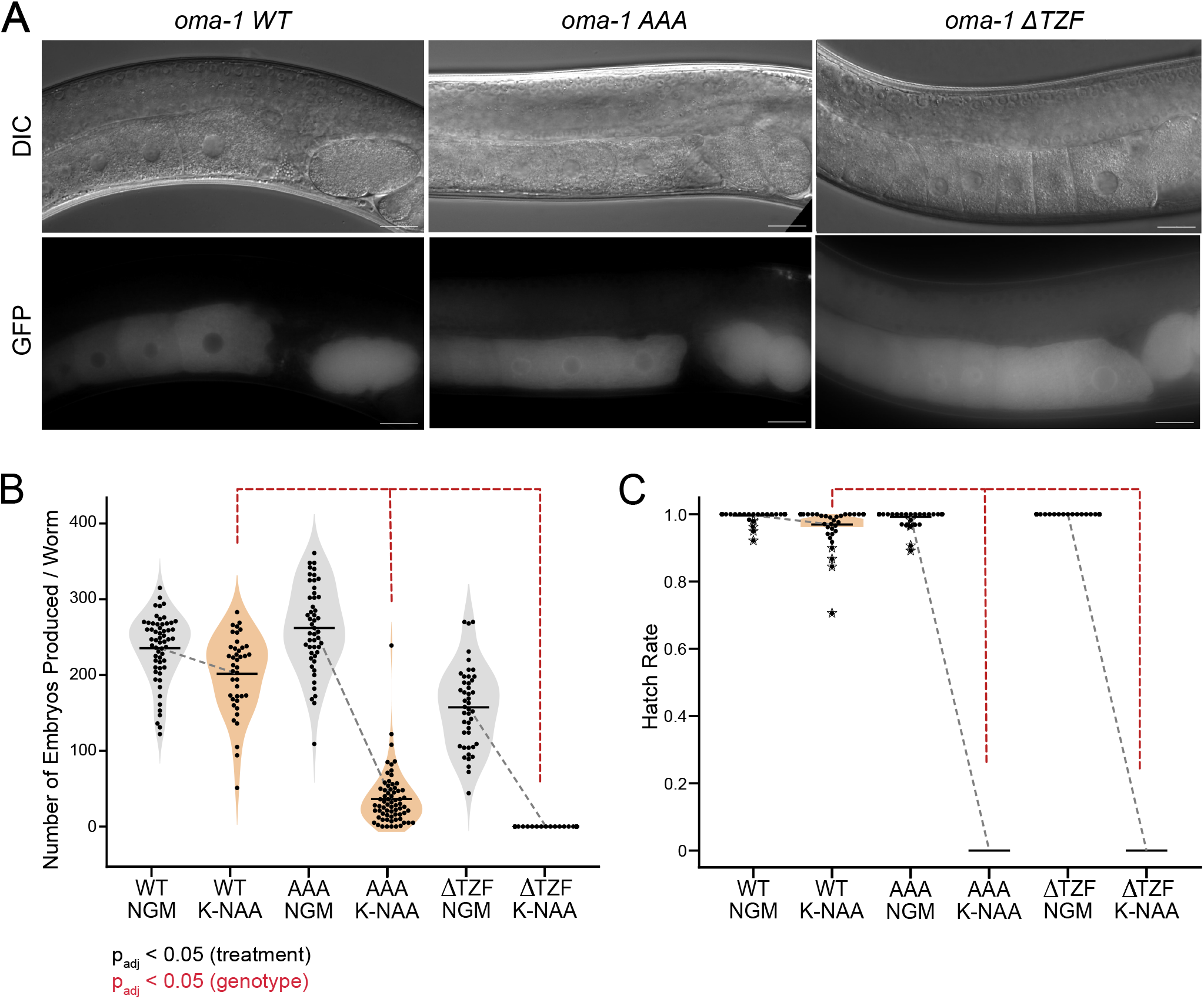
Brood size and hatch rate measurements for *oma-1* variants. **A**. Representative DIC and GFP images of OMA-1::GFP in live worms for the wildtype protein, the RRR to AAA mutant, and the TZF deletion mutant in the absence of K-NAA. The scale bar represents 20 microns. **B**. Total number of embryos produced in all three variants in the absence (gray) and presence (orange) of K-NAA. The individual dots represent the brood of a single adult and the vertical line represents the mean as in figure 3. The statistical significance between conditions (black) and strains (red) is presented as in figure 3. **C**. Hatch rate of embryos produced in all three strains as a function of K-NAA treatment. Dots represent the hatch rate of embryos produced by a single animal. Asterisks indicate outliers identified in Tukey analysis from a one-way ANOVA. Statistical significance is represented as in panel B.

To assess the impact on fertility, we measured the total number of embryos laid, the total number of viable worms produced, and the hatch rate (viable/total) for both genotypes compared to the wild type control cultured under standard growth conditions (NGM-Agar, 20º C, OP50 *Escherichia coli* food, **Fig. 4B–C**). The *oma-1::GFP(ΔTZF)* strain was fertile in the absence of K-NAA (Embryos Produced_ΔTZF, –KNAA_ = 157 ± 53; Viable Brood_ΔTZF, –KNAA_ = 157 ± 53, Hatch Rate = 1.0, N = 45) but completely sterile in the presence of K-NAA (Embryos Produced_ΔTZF, +KNAA_ = 0, Viable Brood_ΔTZF, +KNAA_ = 0, N = 50). By contrast, the control strain produced a similar amount of embryos, most of which remained viable, in the presence or absence of auxin (Embryos Produced_WT, –KNAA_ = 235 ± 44; Viable Brood_WT, –KNAA_ = 235 ± 44, Hatch Rate = 1.0, N = 58; Embryos produced_WT, +KNAA_ = 202 ± 51; Viable Brood_WT, +KNAA_ = 196 ± 51, Hatch Rate = 0.97, N = 41). These data show that loss of the *oma-1* TZF domain in vivo produces an *oma-1* null-like phenotype when *oma-2* is depleted by auxin treatment. As such, the TZF domain is essential *in vivo*.

Next, we measured the total number of embryos produced, the viable brood, and the hatch rate of the *oma-1::GFP(AAA)* mutation (**Fig. 4B-C**). This mutant produced a normal number of embryos in the absence of K-NAA (Embryos Produced_AAA, –KNAA_ = 262 ± 55, Viable Brood _AAA, –KNAA_ = 260 ± 56, Hatch Rate = 1.0, N= 51). However, in the presence of K-NAA, the triple alanine mutation produced a small number of embryos that failed to hatch (Embryos Produced_AAA, +KNAA_ = 33 ± 27, Viable Brood _AAA, –KNAA_ = 0, Hatch Rate = 0, N= 67). The results show that the arginine rich motif upstream from the TZF domain is required to produce viable progeny. However, a small number of embryos were produced in this mutant whereas the ΔTZF domain mutant made none. This suggests that oocyte maturation progresses farther in *oma-1::GFP(AAA)*, but that new problems in fertilized zygotes arise that inhibit successful embryogenesis.

### Mutation of the arginine-rich motif reveals a role for oma-1 in eggshell formation

To better understand the nature of embryonic lethality in the K-NAA-treated *oma-1::GFP(AAA)* mutant worms, we imaged embryos recovered from young adult hermaphrodites. We noted a high preponderance of young adult animals that contained early stage embryos (<100 cells) with polynucleated cells (87%, N=23, **Fig. 5A-B**). In contrast, *oma-1::GFP(AAA)* animals grown in the absence of KNAA did not have polynucleated embryo cells (0%, N=18) and *oma-1::GFP(WT)* animals produced normal embryos both in the absence (N=13) and presence of KNAA (N=10), revealing that the phenotype is unique to *oma-1::GFP(AAA)* animals treated with K-NAA. We note that defects in embryonic divisions have been observed in an *oma-1(te21)*/*oma-2(te50)* double mutant, where both alleles cause incomplete loss of *oma-1/2* function [33].

**Figure 5.**
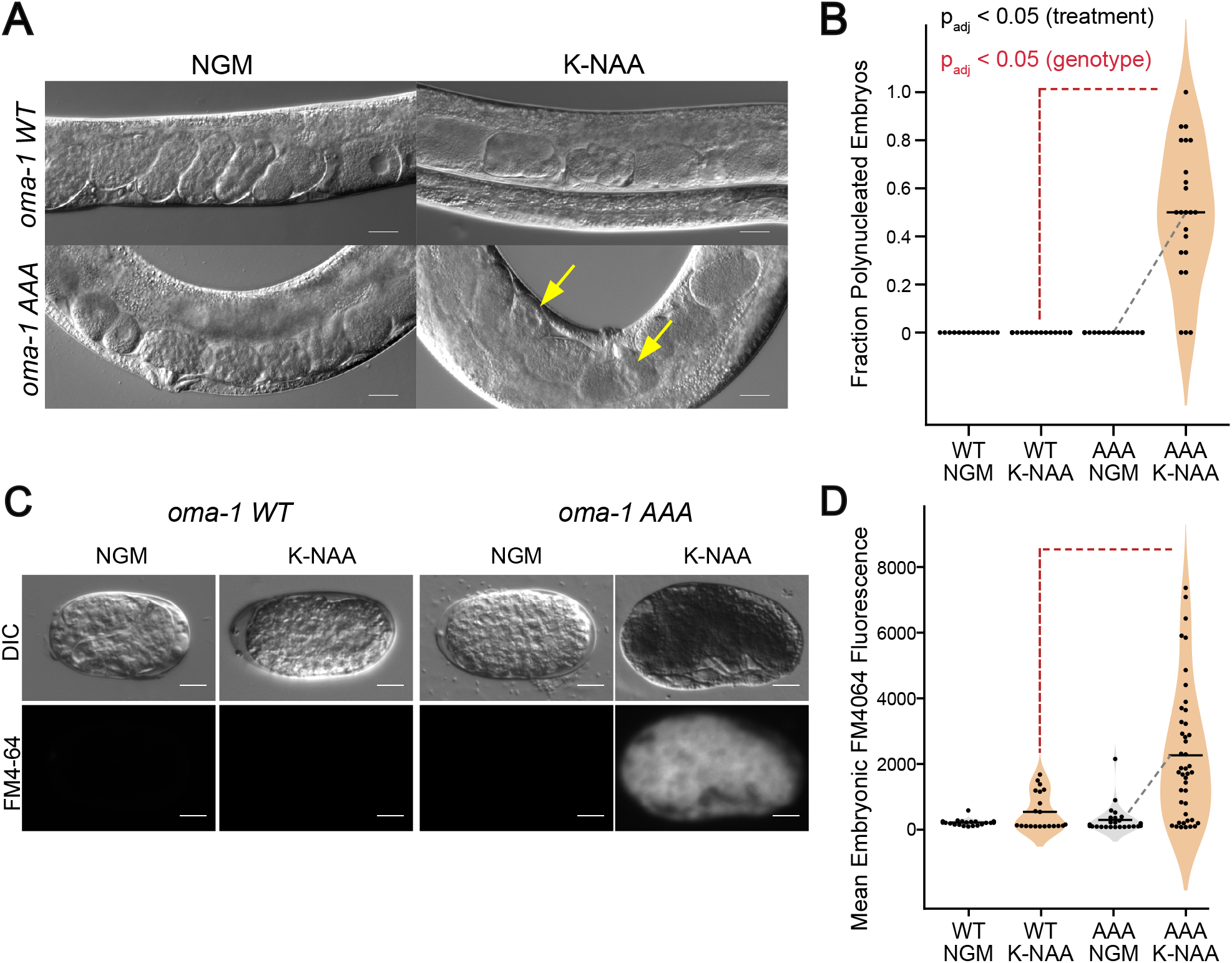
Embryos produced in the *oma-1 (AAA)* mutant have polynucleated embryonic cells and permeable eggshells. **A**. DIC images of WT and AAA mutant animals in the presence and absence of K-NAA. Example poly-nucleated embryos are identified with yellow arrows. The scale bar represents 20 microns. **B**. The fraction of observable embryos with polynucleated cells per adult imaged is shown as a function of genotype and growth condition. The coloring of different samples and the statistical significance are presented as in figure 4. **C**. FM4-64 treatment of embryos recovered by dissection of WT or AAA mutant animals as a function of K-NAA treatment. DIC images and FM4-64 fluorescence images are shown. The scale bar represents 20 microns. **D**. Mean FM4-64 fluorescence intensity in embryos recovered from both strains and both treatment conditions. The coloring and statistical significance are as in figure 4.

Polynucleation of early embryonic blastomeres is a phenotype frequently observed in mutants with defective eggshell production [34]. Nuclei replicate and divide, but mitosis fails at the point of cytokinesis due to weak or incomplete eggshells. To assess the integrity of the eggshell, we performed two experiments. First, we collected embryos from *oma-1::GFP(AAA)* and *oma-1::GFP(WT)* worms, both in the absence and presence of K-NAA, by bleaching. Embryos recovered from KNAA-treated *oma-1::GFP(AAA)* worms did not survive bleach treatment, while untreated *oma-1::GFP(AAA)*, untreated *oma-1::GFP(WT)*, and KNAA treated *oma-1::GFP(WT)* embryos appeared normal and were able to hatch (**Supplemental Figure 3**). These data are consistent with defects in the chitin eggshell, which are typically impervious to strong bleach and base treatment [35].

Next, to quantitatively assess eggshell integrity, we recovered embryos from KNAA-treated and untreated *oma-1::GFP(AAA)* and *oma-1::GFP(WT)* control animals by dissection, which does not require bleach. Recovered embryos were treated with the fluorescent lipophilic dye FM4-64, which cannot transit the *C. elegans* eggshell after it has fully formed (**Fig. 5C-D**) [36]. The mean fluorescence intensity of K-NAA-treated *oma-1::GFP(AAA)* mutant embryos is 9-fold higher than untreated *oma-1::GFP(AAA)* animals (N_KNAA_ = 44, N_ctl_ = 26, P_adj_ = 6.8e-5) and 5-fold higher than KNAA-treated *oma-1::GFP(WT)* animals (N_mut, KNAA_ = 44, N_wt, KNAA_ = 21, P_adj_ = 1.2e-3). The data show that most K-NAA-treated *oma-1::GFP(AAA)* mutant embryos have eggshell integrity defects.

### Embryos break during fertilization

In healthy young adult worms, a mature oocyte is ovulated through the spermatheca, where it is fertilized. Immediately following fertilization, the new zygote emerges into the uterus, where it secretes a hard chitin shell and begins embryonic divisions [2]. To directly observe this process, we filmed ovulation and fertilization in *oma-1::GFP(AAA)* and *oma-1::GFP(WT)* animals both in the presence and absence of K-NAA. Images were collected every two minutes using both DIC and fluorescence optics to visualize ovulation and the pattern of OMA-1::GFP expression.

Surprisingly, we observed the oocyte/zygote fracture in 50% of the movies collected in K-NAA-treated *oma-1::GFP(AAA)* during ovulation (N=8, **Supplemental Movie 1**). In one case, the oocyte fractured into three pieces, with one piece being left behind in gonad, and two pieces remaining in the uterus (**Fig. 6**). We did not observe embryo breaking in the absence of KNAA in *oma-1::GFP(AAA)* movies (N=5) or in presence or absence of K-NAA in *oma-1::GFP(WT)* movies (N=2, N=4 respectively). The data provide further evidence that eggshell formation is aberant in the triple alanine mutant, consistent with a role for OMA-1 in regulating eggshell formation.

**Figure 6.**
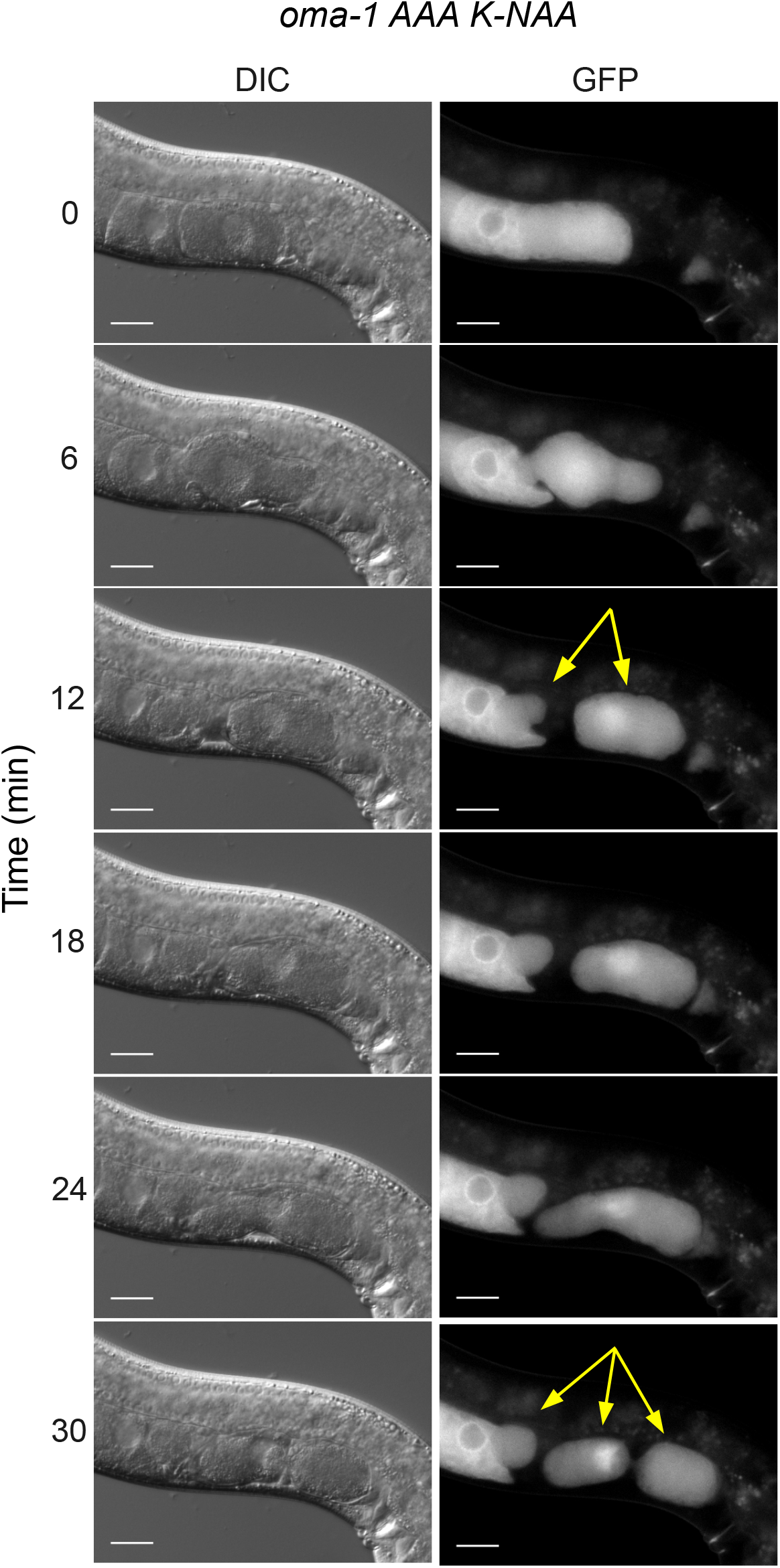
Embryos produced in the *oma-1 AAA* mutant frequently break during ovulation. Still images from a movie of ovulation are shown. DIC and GFP stills are shown. The time post initiation of filming is shown. At the 6 minute time point, and oocyte can be seen transitioning the spermatheca, where it is fertilized. By the 12 minute time point, the new embryo is broken into two pieces, one that remains in the germline, and another in the uterus (yellow arrows). At the 24 minute time point, the embryo in the uterus begins to break again into two pieces to make a total of three embryo fragments (marked with yellow arrows in the 30 minute time point. The scale bars represent 20 microns.

### Numerous gene regulatory pathways are dysregulated in the triple arginine mutation

To gain insight into how the *oma-1::GFP(AAA)* mutant impacts gene expression, we used RNA-seq to measure the transcriptome of mutant and control strains both in the presence and absence of K-NAA. We used DeSeq2 to identify differentially expressed genes between the four groups, comparing data for two biological replicates for each strain at each condition [37].

First, we compared K-NAA-treated *oma-1::GFP(AAA)* to KNAA-treated *oma-1::GFP(WT)* samples (**Fig. 7A-B**). The data reveal 2782 upregulated and 4382 downregulated transcripts caused by the triple arginine mutation with a log_2_ fold-change threshold of 0.585 and a P_adj_ value less than or equal to 0.05. By comparison, untreated *oma-1::GFP(AAA)* vs. untreated *oma-1::GFP(WT)* analyses identified just 138 upregulated and 36 downregulated transcripts, consistent with the presence of functional OMA-2 protein in these animals. 97% of upregulated and 99% of downregulated transcripts are unique to K-NAA-treated *oma-1::GFP(AAA)* samples. We hypothesize that these changes are driven by the mutation and it’s resultant phenotype, rather than arising indirectly from K-NAA exposure.

**Figure 7.**
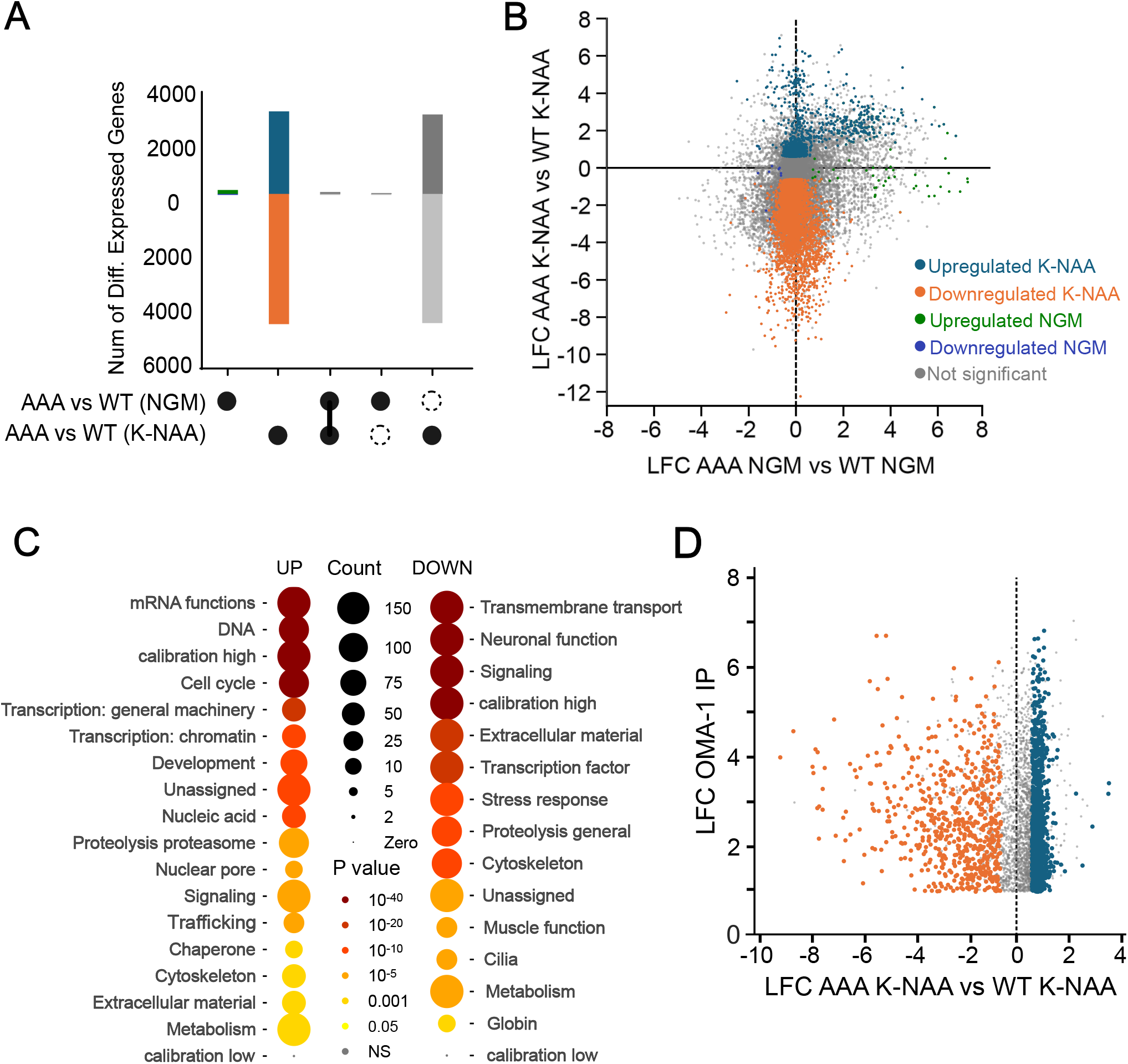
RNAseq analysis comparing gene expression patterns in WT and AAA mutant embryos. **A**. UpSet plots representing the number of genes upregulated (above the 0 line on the y-axis) or downregulated (below the 0 line). The labels under the graph indicate comparisons between gene sets under different conditions. A closed circle indicates the gene sets presented. A vertical line indicates the subset of genes found in both comparisons. A solid circle next to a dashed circle indicates gene sets that are unique to the comparison with the solid circle. The blue color indicates increased genes in the AAA mutant compared to WT controls upon K-NAA treatment. The orange color indicates decreased genes in the same comparison. Green indicates genes increased in the AAA mutant relative to WT under normal growth conditions, while purple represents downregulated genes in the same treatment. **B**. Correlation plot showing that most gene expression changes are unique to the mutant upon K-NAA treatment. The coloring is as in panel A, except here gray represents non-significant changes. **C**. WormCat analysis to identify gene categrories upregulated or downregulated in the mutant compared to control in K-NAA treatment. The size of the circle indicates the number of genes in the set, the color indicates statistical significance. **D**. Comparison of the LFC of mRNAs enriched in an OMA-1 IP [18] compared to the genes that change in in the AAA mutant relative to control upon K-NAA treatment. The coloring is as in panel A.

To better understand which sets of genes are dysregulated in the triple alanine mutation, we performed WormCat analysis on the genes that were differentially expressed comparing KNAA-treated *oma-1::GFP(AAA)* to KNAA-treated *oma-1::GFP(WT)* (**Fig. 7C**) [38]. The results show numerous upregulated and downregulated gene categories. The mRNA function, DNA, and cell cycle gene categories are the most signficantly upregulated, consistent with a role for OMA-1 in regulating the oocyte-to-embryo transition and controlling the expression of maternally-supplied mRNAs [7, 17, 18]. Transmembrane transport, neuronal function, and signalling gene categories were the most significantly downregulated. Transport and signalling are involved in oocyte maturation and cell fate specification in embryos [2].

### OMA-1-associated mRNAs are dysregulated in the triple alanine mutant

Next, we looked to see if known direct targets of OMA-1 are dysregulated in our data sets. OMA-1 has been shown to regulate *glp-1, zif-1, mei-1, nos-2, mex-3, atg-4*.*2, cul-4, ets-4*, and *set-6* through interactions with their 3’UTR [2, 19-21]. Of these, six are upregulated in K-NAA-treated *oma-1::GFP(AAA)* samples compared to K-NAA treated WT controls (*glp-1, zif-1, mei-1, nos-2, mex-3*, and *cul-4*). One is downregualted in the mutant compared to control (*ets-4*), and two are unchanged (*atg-4*.*2, set-6*) (**Supplemental Data Set 2**). As such, 78% of transcripts where gene-specific studies have been carried out are dysregulated in the *oma-1::GFP(AAA)* mutant. Next, we compared a list of OMA-1 bound mRNAs to our RNA-seq data. Greenstein and colleagues identified OMA-1 interacting mRNAs using immunoprecipitation and RNA sequencing [18]. Of the OMA-1-associating mRNAs enriched at least two fold over controls in immunoprecipitations, almost half (47% 1608/3414) are dysregulated in *oma-1::GFP(AAA)* samples compared to WT controls (**Fig. 7D**). Of these, 29% (973/3414) are upregulated, and 19% (635/3414) are downregulated.

### Several genes essential to eggshell formation are dysregulated in the triple arginine mutant

To identify genes that may be involved in the fragile eggshell and cytokinesis phenotypes observed in the triple arginine mutant, we compared the list of dysregulated transcript in our mutant to all *C. elegans* genes that include the terms “eggshell” or “fertilization defective” in the phenotype field listed in WormBase [39]. We identified 37 genes in our RNA-seq data that intersect with these phenotype categories. Of these, 42% (15/37) are dysregulated in the triple alanine mutant, including 31% that are upregulated (11/37) and 11% that are downregulated (4/37).

Intriguingly, the upregulated genes include *egg-1, egg-2*, and *egg-3*. EGG-1 and EGG-2 are redundant oocyte membrane-localized LDL-receptor proteins necessary for localization of chitin synthase, eggshell formation, and the block to polyspermy [40, 41]. EGG-3 is a protein tyrosine kinase that, along with MBK-2, is also necessary for chitin synthase localization and eggshell formation [42, 43]. The *sep-1* gene, a cysteine protease, is also upregulated in the triple alanine mutant. SEP-1 is essential to several early embryonic cell division events, including sister chromatid separation, eggshell formation, and polar body extrusion [44].

In contrast, of the four downregulated genes, only *kgb-1* appears to play a specific role in fertilization. This gene encodes a serine/threonine kinase in the JUN kinase (JNK) family. It has been shown to associate with all four germline helicase proteins that associate with p-granules (glh-1/2/3/4), and it plays a role in oocyte maturation [45]. We suggest that these dysregulated genes contribute to the eggshell and cytokinesis defects we observe in the *oma-1::GFP(AAA)* mutant.

## DISCUSSION

In this study, we show that a thirteen amino acid arginine rich domain upstream of the OMA-1 tandem zinc finger domain is essential for OMA-1 RNA-binding activity. Unlike other members of this family, the TZF domain is not sufficient for high affinity binding. Importantly, this region contributes to both RNA-binding affinity and positive cooperativity—another property that seems to be unique to OMA-1 compared to other TZF domain proteins.

We also show that three adjacent arginine residues in this peptide are essential for fertility in vivo. However, unlike a mutation that removes the TZF domain, the triple arginine to alanine mutation can be fertilized and produce embryos. These embryos have weak eggshells, frequently break, and arrest prior to morphogenesis, demonstrating that this region is essential for OMA-1 function in vivo. However, because fertilization and early embryogenesis proceeds further in the *oma-1::GFP(AAA)* animals than in *oma-1::GFP(ΔTZF)* mutants or other nulls, we suspect that it acts like a genetic hypomorph, retaining some OMA-1 activity, but not enough to successfully navigate early embryogenesis.

### Why is the TZF domain insufficient for RNA-binding activity?

Positively charged residues, arginines and lysines, play a crucial role in protein-RNA interactions [26, 29]. A statistical analysis of protein-RNA complexes indicated that lysine and arginine residues have the highest propensity to be found at protein-RNA interfaces, with Arg being the most dominant [29]. Both residues contribute to the stability of the complex through electrostatic interactions with the RNA backbone phosphate groups due to their positive charge. In addition, they can facilitate nucleic acid recognition by transiently forming non-specific electrostatic interactions in the encounter complex, thereby reducing the search from a three-dimensional to a one-dimensional space [46, 47].

Previous studies have shown that TZF domains of various ZFP36-like CCCH-type tandem zinc finger proteins (TIS11d, TTP, MEX-5 and POS-1), are necessary and sufficient for binding their target RNAs [12-14]. Under physiological conditions, the TZF domains of these proteins are highly positively charged (**Supplemental Figure 1**). In contrast, the TZF domain OMA-1 contains fewer positively charged residues and a higher proportion of negatively charged residues compared to the other family members. More specifically, the N-terminal zinc finger of OMA-1 contains three positively charged and four negatively charged residues, resulting in a net negative charge.

Based on our binding data, we postulate that the arginine-rich region upstream of zinc finger 1 evolved to compensate for the reduced number of positively charged residues within zinc finger 1. This highly positively charged region may contribute to RNA recognition by mediating non-specific electrostatic interactions, either during the formation of the initial recognition complex or by stabilizing the protein-RNA interface through interactions with the RNA backbone. The similar binding affinities observed for the wildtype OMA-1 (RRR) and the KKK mutant support our hypothesis that these interactions are electrostatic and non-sequence specific.

### How do the RNA-binding properties of OMA-1 contribute to its function in worms?

It is clear from our binding data that OMA-1(1-182)-AAA binds to RNA with weaker apparent affinity and reduced cooperativity when measured against a variety of target mRNA sequences. How might reduced affinity and cooperativity impact OMA-1 function? We do not know the absolute concentration of OMA-1 in oocytes and embryos, but we do know that the relative OMA-1 concentration increases in maturing oocytes, then decreases quickly after the first cell division. The triple arginine mutation does not appear to impact the relative abundance or dynamics of OMA-1::GFP expression in vivo. As such, it is possible that the phenotypes we observe are directly caused by the reduction in apparent binding affinity and cooperativity. For example, some RNAs with multiple OMA-1 binding motifs (OBMs) might still be repressed by a weakened version of OMA-1 due to statistical effects, while other RNAs with fewer binding sites might be dysregulated. This might account for the ability of mutant oocytes to be fertilized yet fail at a later stage during embryogenesis. However, we cannot rule out the possibility that the three arginine residues perform other roles in vivo, including binding to unidentified regulatory co-factors or repressive machinery, or possibly through modification by arginine methylases, or similar. Therefore, though reduced affinity provides a plausible explanation for the phenotypes we observe, more work will be needed to define the precise mechanism by which these amino acids contribute to OMA-1 function.

### What is the nature of embryonic lethality in the triple arginine mutant?

The most striking features of the *oma-1::GFP(AAA)* mutant are the weak eggshells, polynucleated embryonic cells, and frequent breakage during fertilization. These phenotypes are likely connected. It has long been known that the eggshell contributes to cytokinesis, and mutants that impact proteoglycan synthesis, including *sqv-6, sqv-2*, and others often form polynucleated embryos. Our RNA-sequencing data revealed many genes whose abundance changes as a function of the triple arginine mutation, but five of these (*egg-1, egg-2, egg-3, sep-1, and kgb-1*) have well-characterized roles in eggshell formation. All of these have multiple OBMs in their 3’UTRs and may be direct regulatory targets of OMA-1. It remains possible that other genes that are dysregulated could contribute, as there are thousands. OMA-1 has been proposed to regulate the translation of several maternal genes, but a role in the regulation of mRNA stability has not been ruled out. More work will be needed to dissect the molecular mechanisms by which OMA-1 regulates its mRNA targets, and whether the phenotypes observed upon *oma-1::GFP(ΔTZF)* or *oma-1::GFP(AAA)* can be explained by the dysregulation of one or a few key target mRNAs, or if the phenotypes are caused by the aggregate results of dysregulating many genes.

Our work shows how a structure-to-phenotype approach, where quantitative in vitro studies guide in vivo mutagenesis, can yield new insights into the role of even well characterized genes in fundamental biological processes.

## MATERIALS AND METHODS

### Protein purification

The genes corresponding to residues 1-182, 98-182 and 110-182 were amplified and subcloned into petSUMO vectors (pet28a with N-terminal hexa-his-SUMO fusion tag). All plasmids were transformed into BL21 (DE3) *Escherichia coli* cells. The cells were cultured at 37°C in LB broth for binding studies, and in M9 minimal medium for NMR experiments. M9 minimal medium was supplemented with ^15^NH_4_Cl as the sole nitrogen source to achieve uniform ^15^N-labeling. When the OD_600_ reaches ∼0.6-0.8, the culture was supplemented with 100 μM Zn(OAc)_2_, and protein expression was induced by adding isopropyl-β-D-thiogalactopyranoside (IPTG) to a final concentration of 1mM. The culture was incubated overnight at 20°C. Cells were then harvested by centrifugation, and the resulting pellets were stored at -20°C until use.

When ready to use, the cell pellets were resuspended in 50 mM Tris, 300 mM NaCl, 100 μM Zn(OAc)_2_, 100 μM TCEP and 25 mM imidazole and lysed by sonication. The lysate was centrifuged to separate soluble fraction from insoluble fraction at 19,000 rpm (∼45,000 g) for 45 min at 4°C. The supernatant was loaded onto Ni-NTA column (HisTrap HP, Cytiva), washed with the same buffer, and finally the protein was eluted via gradient of imidazole concentration from 25 mM to 300 mM. Elution fractions containing SUMO-tagged OMA-1 were pooled and dialyzed into a buffer containing 50 mM Tris, 50 mM NaCl, 100 μM Zn(OAc)_2_, 100 μM TCEP. Ubiquitin-like specific protease 1 (Ulp1) was added to the sample to remove the SUMO-tag during overnight dialysis. The protease and SUMO-tag were removed by ion exchange chromatography. Finally, the protein sample was further purified using size exclusion column equilibrated with a buffer containing 50 mM MES pH 6.0, 100 mM NaCl, 100 μM TCEP and 100 μM Zn(OAc)_2_. For purification of the OMA-1(1-182) construct, 50 mM arginine and 50 mM glutamate were added to all buffers to reduce aggregation.

OMA-1 cloned into the pHMTc vector was used for binding studies of the OMA-1 variants. This vector encodes an OMA-1 N-terminally tagged with hexa-His tag and maltose binding protein (MBP), which enhances the solubility and the stability of the protein. The AAA and KKK mutants were generated using the Quick-Change mutagenesis protocol. Protein expression and purification were carried out following a similar procedure to that described above. Following the Ni-NTA affinity chromatography, the eluate was loaded onto amylose resin (NEB), which has high affinity for MBP. After washing, the MBP-tagged protein was eluted with buffer containing 5mM maltose (50 mM Tris, 100 mM NaCl, 100 μM Zn(OAc)_2_, 100 μM TCEP). The eluate from amylose resin was further purified by ion exchange chromatography.

At every step of the purification, the purity of the sample was confirmed by SDS-PAGE gel. The protein concentration of the samples was estimated from UV-absorption at 280 nm, using a nanodrop (ThermoFisher Scientific).

### RNA-binding assay

The binding affinity of the OMA-1 constructs to various RNA sequences was carried out by Electrophoretic Mobility shift Assay (EMSA). In this assay, OMA-1 at different concentrations was incubated with fluorescein labeled RNA (IDT) constructs in 50 mM Tris, pH 8.0, 100 mM KCl, 100 μM Zn(OAc)_2_, 0.1 mg/mL tRNA, 0.01% IGEPAL, and 1 mM DTT at room temperature for 3hrs to reach equilibration. The equilibrated samples, then, were loaded onto 8% acrylamide gels prepared with TB buffer (89 mM Tris and 89 mM Boric acid). The gel was exposed to 120V electric potential for 90 min at 4°C to allow the free and bound RNA chains migrate through the gel.

To detect the fluorescently labeled RNA bands, the gel was imaged using Fuji FLA-5000 Fluoro Image Analyzer and then the band intensities were estimated using ImageJ. The fraction of bound RNA was calculated from the intensities of the bands corresponding to free and bound states for each sample. Then the apparent dissociation constant, K_d,app_, was determined by fitting the fraction of bound RNA at increasing OMA-1 concentrations using the quadratic equation:

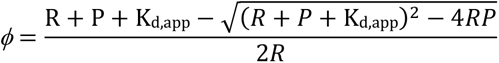

### NMR spectroscopic studies

For NMR spectroscopy, OMA-1(98-182) and OMA-1(110-182) samples were dialyzed into a buffer containing 50 mM MOPS, pH 6.0, 300 mM NaCl, 100 μM TCEP, 100 μM Zn(OAc)_2_, 5% D_2_O was added before data collection. For both samples, ^15^N^1^H-HSQC spectra were collected on a 600 MHz Varian Inova spectrometer. The raw FID data was processed using NMRPipe[48] and visualized using Sparky[49].

### Strains and nematode culture

All strains used in this study are listed in **Table 1**. Strains were maintained by growing animals on NGM (Nematode Growth Medium) seeded with *E. coli* (OP50) under standard conditions [50]. All primers used to produce or evaluate the strains are listed in **Supplemental Table 1**. Strains expressing *oma-1::GFP* (IV) and *deg::oma-2*;*Tir1::mRuby* were gifts from Craig Mello’s lab (UMass Chan Medical School). WRM66 was made by crossing male *oma-1::GFP* animals with hermaphrodite *AID::oma-2*;*Tir1::mRuby* animals.

**Table 1.**

| Strain ID | Genotype |
| --- | --- |
| WRM66 | *oma-1::GFP* IV, *deg::oma-2*, *Tir1::mRuby* V |
| WRM92 | *oma-1(spr24[RRR-AAA])::GFP* IV, *deg::oma-2*, *Tir1::mRuby* V |
| COP2772 | *oma-1(knu1284[P110-H182del])::GFP IV, deg::oma-2, Tir1::mRuby V* |

The RRR-to-AAA mutant (WRM92) was generated in the WRM66 strain background using CRISPR-Cas9 genome editing following the procedure of Ghanta and Mello [51]. To produced WRM92, WRM66 worms were injected with an RNP mixture containing *Streptococcus pyogenes* Cas9, a crRNA (sgRNA_92), tracrRNA, and an alt-R modified homology-directed repair donor oligo (Oma-1_AAA). The pRF::rol-6 plasmid was included in the mix as an injection control [51]. The mutation recodes nine nucleotides that are 30-21 base pairs upstream of the TZF domain from AGA-CGC-CGC (RRR) to GCT-GCT-GCA (AAA). The presence of the desired mutation was detected by PstI restriction digestion of a PCR product (oma1_seq_F1 and oma1_seq_R2). The mutation was confirmed by Sanger sequencing.

The TZF knockout mutant (COP2772) was generated using two sgRNAs (sgRNA_94a and sgRNA_94b), and an oligonucleotide donor homology guide DNA (pNU3644odn) in the background strain WRM66. The mutation removes 261 base pairs in the *oma-1* coding sequence and recodes 19 base pairs in exon 3 with silent mutations to prevent re-cutting of the sgRNAs. The strain was validated with PCR and sequencing. This strain was generated and validated by InVivo Biosystems (Eugene, OR).

### Brood size and hatch rate assays

Worms were synchronized by bleaching with a 20% bleach solution (3 ml concentrated bleach, 3.75 ml 1M filtered sodium hydroxide, 8.75 ml H2O), then washing the remaining embryos with M9 twice before allowing them to hatch in the M9 overnight. One day after bleaching, L1 worms were plated onto NGM agar plates seeded with OP50 and left to grow until the L4 larval stage, after which the worms were individually transferred onto either NGM plates or NGM plates containing 4mM K-NAA. For the experiments shown in Figure 3, 1 mM IPTG and 100 mg/mL Ampicillin were also added to the agar plates. For each day up to six days post hatch, the individual worms were transferred to new plates, and the remaining embryos on the previous plate were counted. The number of viable progeny was counted from the embryos that were laid in the previous day. Embryos or hatchlings found on the plate seven days post hatch were included in the overall count, but the parent worms were not transferred. At the end of the experiment, all embryo and hatchling counts were added up to get the total number of embryos and viable progeny each worm produced. The total hatch rate was calculated by dividing the sum of viable progeny by the sum of total embryos laid. Brood size and hatch rate assays on both standard and auxin plates were done for all strains analyzed in this study.

### Nematode imaging and phenotyping

Young adult worms were immobilized with 5-7 microliters of 1 mM levamisole and mounted on 2% agarose pads prior to imaging on a Zeiss AxioObserver 7 at 200X total magnification with both DIC and fluorescnece optics. Embryos were imaged in utero. The number of polynucleated embryos (embryos that contained at least one cell with more than two nuclei) in a single worm was divided by the total number of embryos in the worm to obtain the fraction of polynucleated embryos per animal.

In the FM4-64 dye permeability experiments, embryos were from adults via dissection, then washed in M9, recovered by centrifugation, then washed again with M9 prior to treatment. Embryos were soaked in M9 containing 1 mM FM4-64 for 2-3 hours at 20 degrees C. The embryos were then washed twice with M9 again to remove excess dye and a small amount of the remaining pellet (3-5 microliters) was pipetted onto a 2% agarose pad for imaging at 400X with DIC and fluorescence optics.

### Embryogenesis movies

Young adult worms were harvested from NGM or NGM K-NAA plates, immobilized with 25 mM levamisole, and mounted on 2% agarose for imaging as described in the previous section. DIC and GFP images were collected every two minutes at 200X total magnification for a total time of 50 minutes. Focus was maintained throughout imaging with a Zeiss Definite Focus unit. The embryos were scored for breakage in the movies independently by two lab members.

### RNA sequencing

Prior to RNA isolation, WT and AAA mutant worms were synchronized as described previously with bleach and plated onto NGM plates with or without 4 mM K-NAA. Two 60 mm plates for each strain/condition combination were cultured. Four days post hatch, the worms were harvested from the plates and washed at least three times with nuclease free water to remove OP50. RNA was isolated from each sample using Trizol, chloroform, and isopropanol. Ribosomal RNA from each sample was depleted using a rRNA depletion protocol for *C. elegans* as detailed in Duan et al, 2020 [52]. Library prep was performed with the NEB-Next Ultra II library prep kit (E7775S) following the recommended protocol. Library indexing was done with oligos from the NEBNext Multiplex Oligos for Illumina (Dual Index Primer Set 1 and 2: Cat#: E7335S, E7500S). The concentration for each library was determined by both Qubit and fragment bioanalyzer. Barcoded libraries were sequenced on an Illumina NextSeq1000 with a P1 100 cycle flow cell.

The resulting raw sequencing reads (formatted as FASTQ files) were analyzed using the OneStopRNAseq pipeline [53]. FastQC. v.0.11.5 and MultiQC were used for raw and post-alignment quality control, respectively. STAR v2.7.5a was used to align the reads to the reference genome using WBcel235.90 annotations with default settings except for the following parameters ‘-Q 20 –minOverlap 1 --fracOverlap 0 -p -B -C’ for paired-end strict-mode analysis [54]. Differential expression (DE) analysis was performed with DESeq2 v1.28.1 [37]. Significantly differentially expressed genes were filtered with the criteria FDR < 0.05 and absolute log2 fold change (|LFC|) > 0.585.

## Supporting information

Supplemental Data Set 1

Supplemental Data Set 2

## Data availability statement

The data and statistical analyses presented in in figure 3 through figure 5 of this work are available in **Supplemental Data Sets 1** and **2**. All sequencing data have been uploaded to the NCBI Sequence Read Archive under Bioproject PRJNA1260521.

## ACKNOWLEDGEMENTS

The authors thank Dr. Krishna Ghanta and Dr. Ebru Kaymak for technical support and helpful discussions concerning strain generation and characterization. We thank Dr. Yekaterina Makeyeva and Dr. Craig Mello for strains harboring *oma-1::gfp, deg::oma-2*, and *Tir1::mRuby* alleles used to make the strains described here. We thank Dr. Ye Duan for technical support with the rRNA depletion protocol. This work was supported by NIH Grants R01GM139316 to F.M. and S.P.R. and R01GM145062 to S.P.R.

## AUTHOR CONTRIBUTIONS

AE performed and analyzed the experiments performed in Figures 1 and 2, including all binding experiments and NMR studies. SR generated the WRM66 background strain presented in Figure 3. SN generated and validated the *oma-1(AAA)* mutant strain. SN also collected and analyzed the data presented in Figures 3–6, except for the ΔTZF data presented in Figure 4, which was collected by CD. SN collected the data presented in figure 7, which were analyzed by SN and SR. AE, SN, FM, and SR wrote the manuscript, which was read and approved by all authors.

## SUPPLEMENTARY FIGURE LEGENDS

**Supplemental Figure 1.**
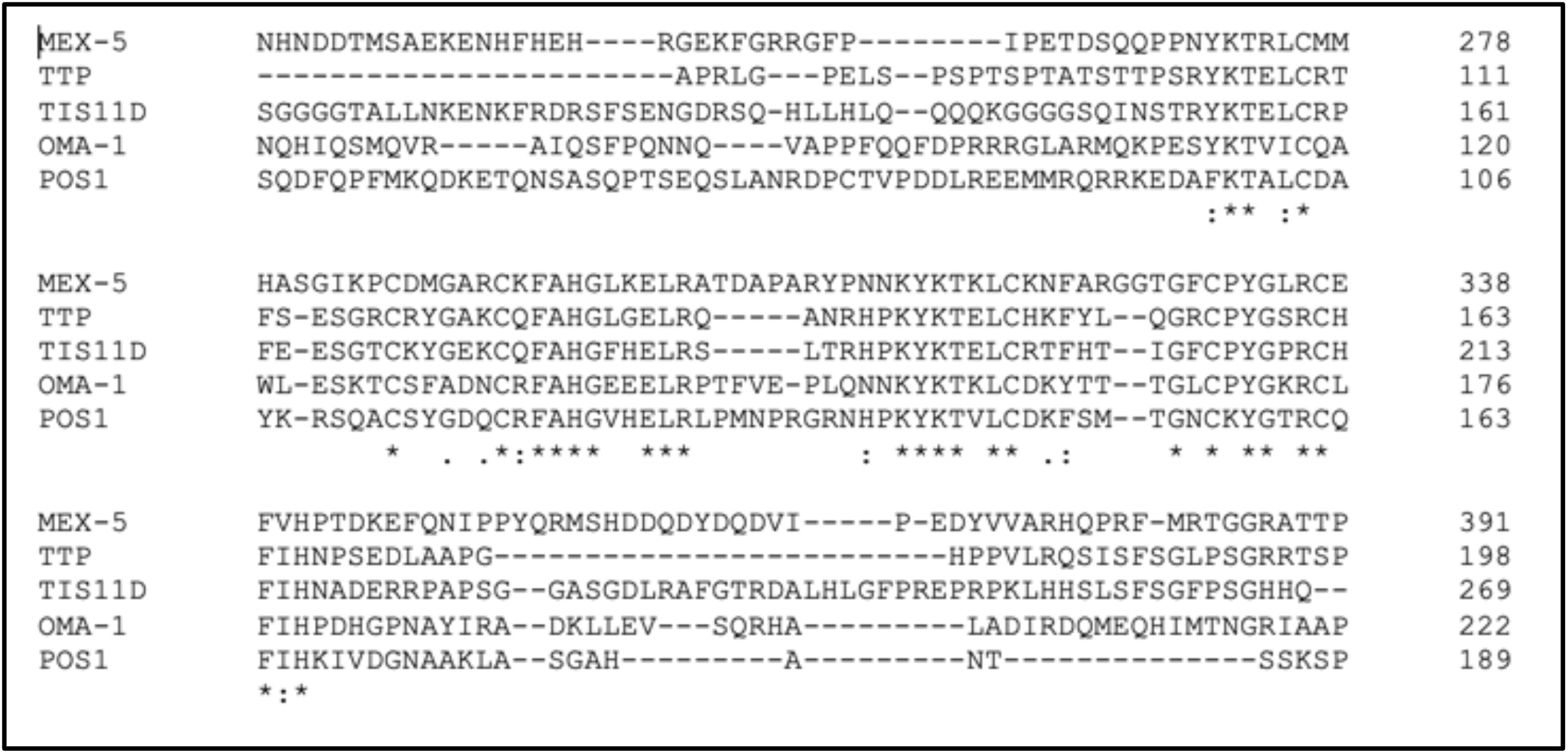
Multiple sequence alignment of the TZF region from a subset of CCCH-type tandem zinc finger domain proteins from vertebrates (TTP, TIS11D) and nematodes (MEX-5, OMA-1, POS-1). Asterisks indicate conserved amino acids, colons indicate amino acid similarity.

**Supplemental Figure 2.**
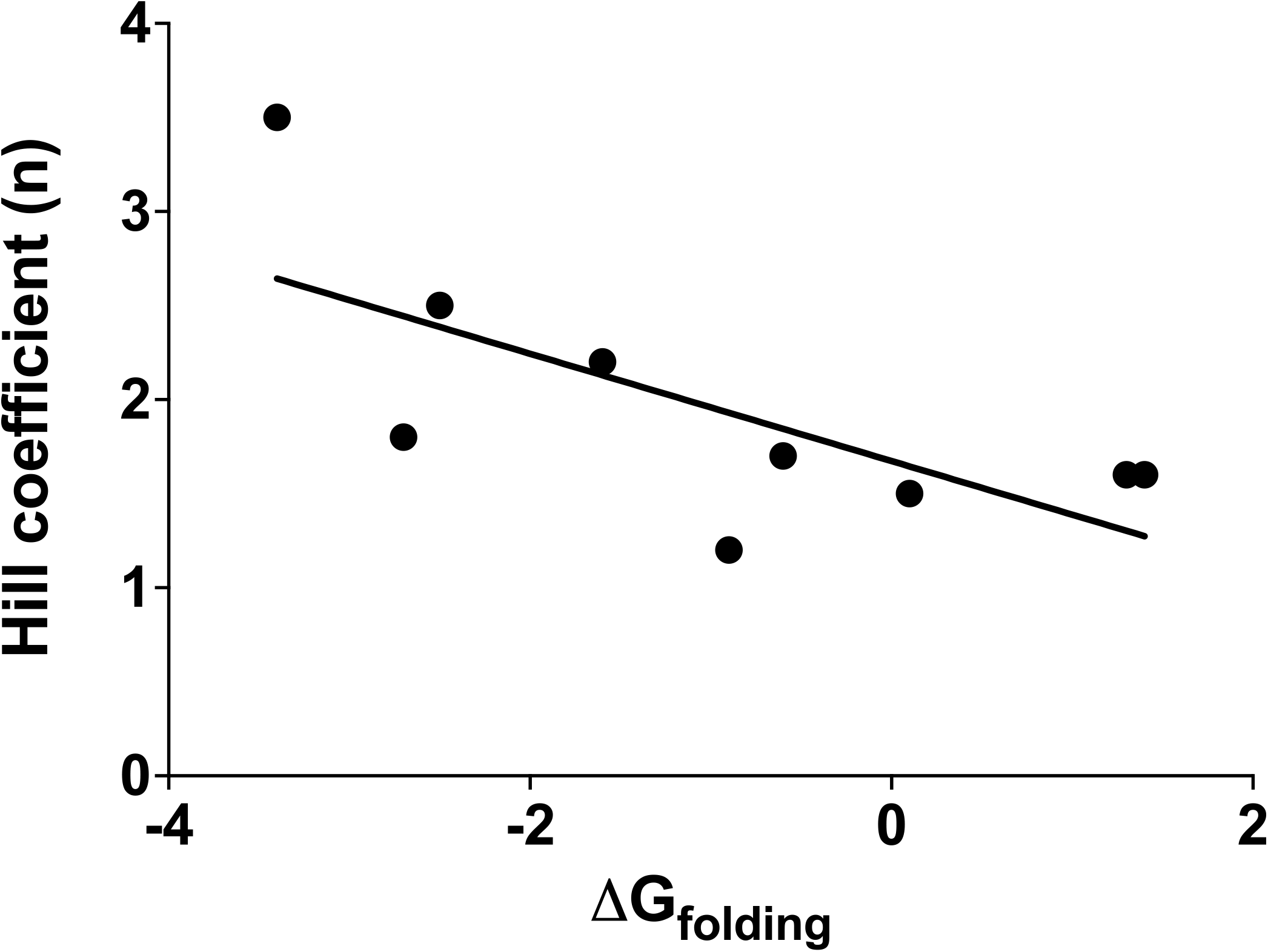
The calculated free energy of RNA folding for sequences that associate with OMA-1 is anticorrelated with the measured Hill coefficient [17, 55]. Each dot represents a different RNA sequence. The line represents a linear fit to the data.

**Supplemental Figure 3.**
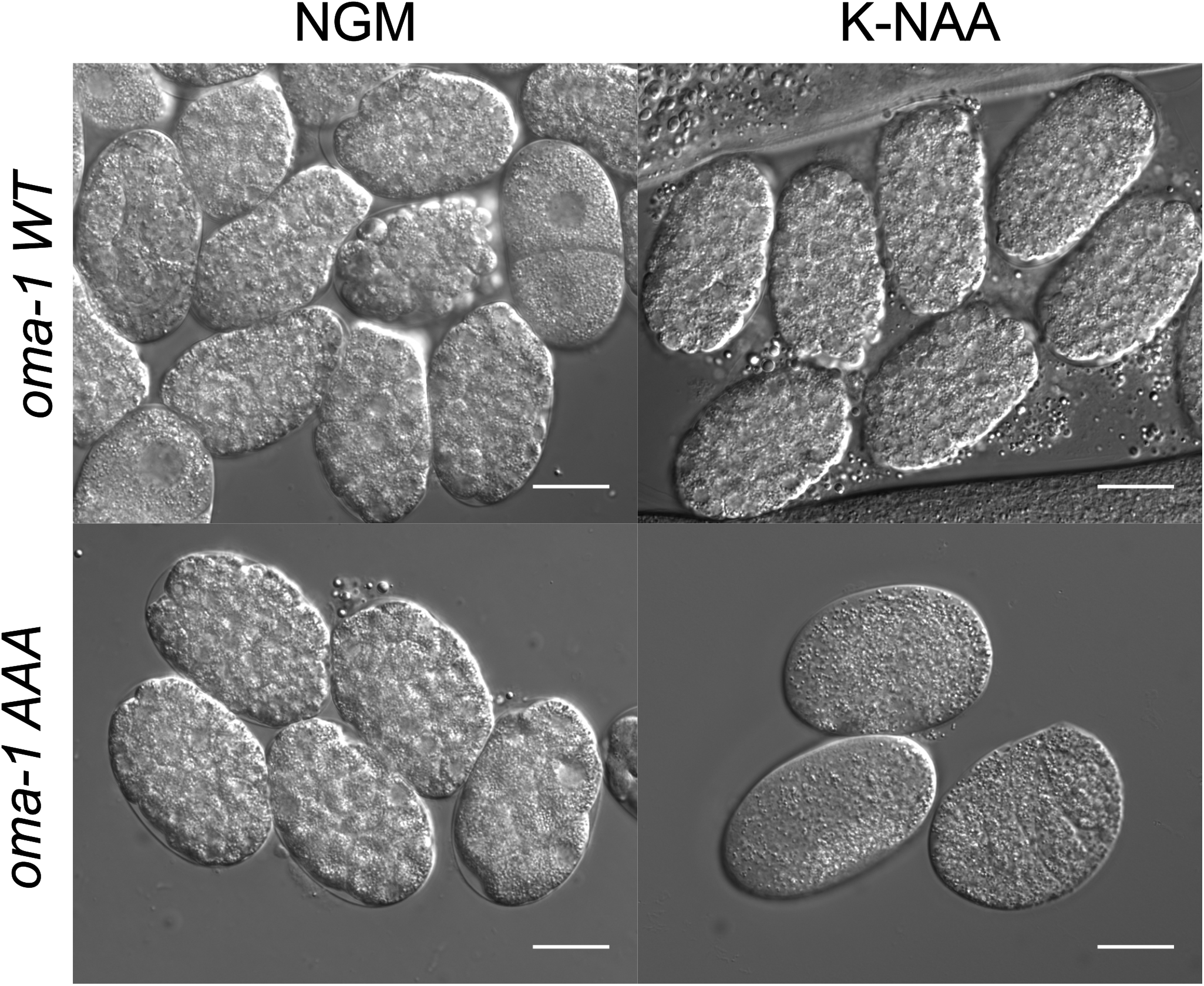
Sample images comparing the state of bleached embryos recovered from *oma-1::GFP* (WT) and *oma-1::GFP* (AAA) mutant worms grown in the presence and absence of K-NAA. The scale bar represents 20 microns. Images were collected using a DIC optics at 400x total magnification.

## SUPPLEMENTARY MOVIE LEGENDS

**Supplemental Movie 1** Example DIC (left) and GFP (right) movies are shown for an *oma-1::GFP (AAA)* animal in the presence of K-NAA (auxin). The time course in minutes is shown on the top left of each panel. The scale bar represents 20 microns.

**Supplemental Movie 2** Example DIC (left) and GFP (right) movies are shown for an *oma-1::GFP (AAA)* animal in the absence of K-NAA (auxin). The scale and time course are as in Supplemental Movie 1.

## SUPPLEMENTARY FILES

1. Supplemental Figure 1: Multiple sequence alignment of the TZF regions.
2. Supplemental Table 1: List of primers used in the study
3. Supplemental Data Set 1: Excel spreadsheet containing all data presented in the figures 1-5.
4. Supplemental Data Set 2: Excel spreadsheet of analyzed RNA-seq data in figure 7.
5. Supplemental Movie 1: Ovulation movie with K-NAA.
6. Supplemental Movie 2: Ovulation movie with standard NGM.

## SUPPLEMENTARY TABLE

**Supplemental Table 1.**
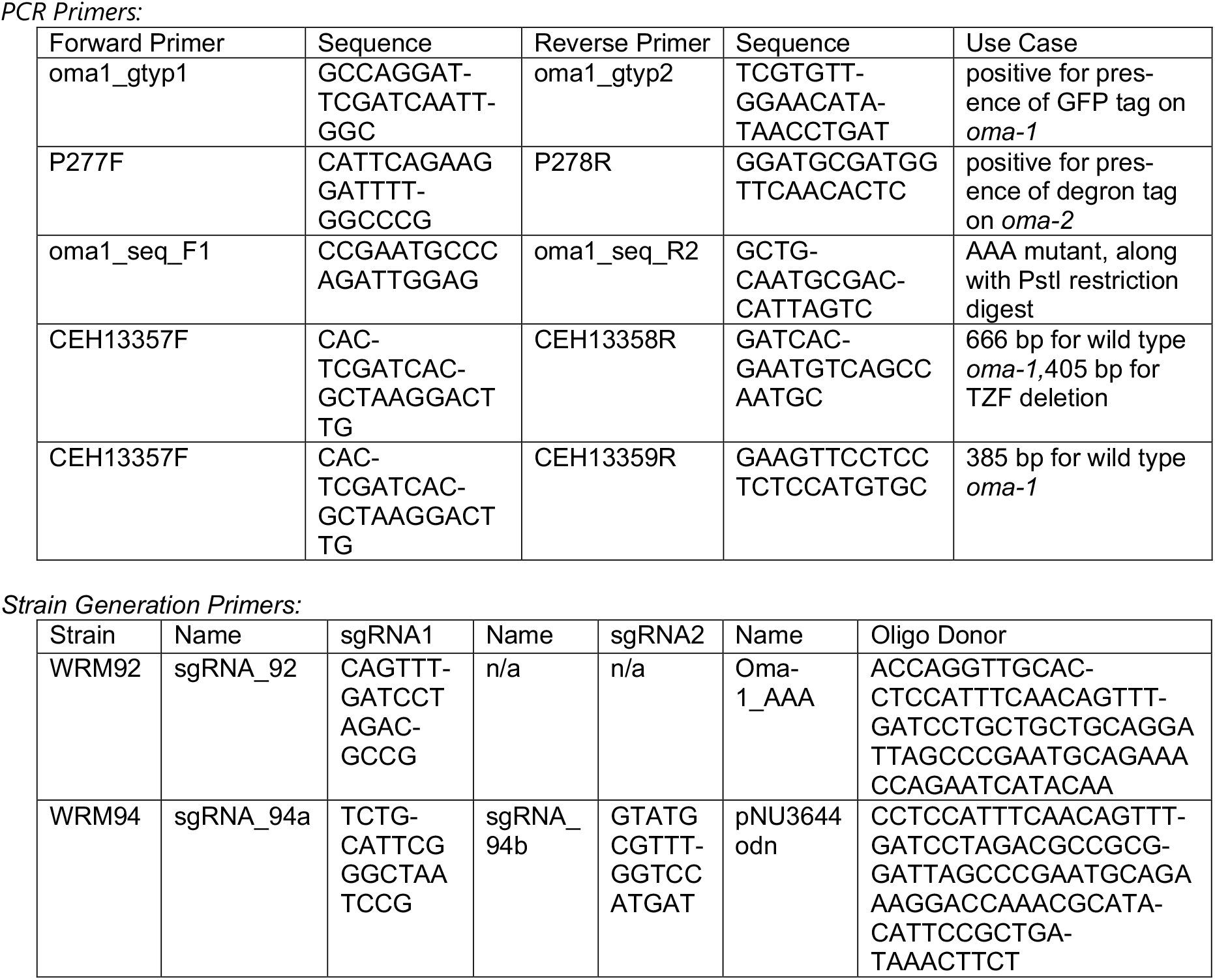

